# Imbalanced Unfolded Protein Response Signaling Contributes to 1-Deoxysphingolipid Retinal Toxicity

**DOI:** 10.1101/2022.09.22.509071

**Authors:** Jessica D. Rosarda, Sarah Giles, Sarah Harkins-Perry, Elizabeth A Mills, Martin Friedlander, R. Luke Wiseman, Kevin T. Eade

**Affiliations:** Department of Molecular Medicine, The Scripps Research Institute, La Jolla, CA 92037; Lowy Medical Research Institute, La Jolla, CA 92037

**Keywords:** deoxysphingolipid, unfolded protein response, retinal degeneration, ATF6, PERK, photoreceptor, retinal organoids, retina

## Abstract

1-Deoxysphingolipids (1-dSLs) are atypical cytotoxic sphingolipids formed through the substitution of alanine for serine in de novo sphingolipid biosynthesis. Accumulation of 1-dSLs has been linked to diseases of the eye such as diabetic retinopathy and Macular Telangiectasia Type 2 (MacTel). However, the molecular mechanisms by which 1-dSLs induce toxicity in retinal cells remains poorly understood. Here, we integrate bulk and single-nucleus RNA-sequencing to define the biological pathways that contribute to toxicity caused by the 1-dSL species, 1-deoxysphinganine (1-dSA), in human retinal organoids. Our results demonstrate that 1-dSA preferentially and differentially activates signaling arms of the unfolded protein response (UPR) in photoreceptor cells and Müller glia within retinal organoids. Using a combination of pharmacologic inhibitors and activators, we define the roles for individual arms of the UPR in 1-dSL-mediated toxicity. We show that sustained PERK signaling through the integrated stress response (ISR) promotes 1-dSL-induced apoptosis in photoreceptors. In contrast, deficiencies in signaling through the ATF6 arm of the UPR contribute to photoreceptor toxicity. These results indicate that imbalanced signaling between the pro-apoptotic PERK/ISR and protective ATF6 arms of the UPR contributes to 1-dSL-induced photoreceptor toxicity. Further, our results identify new opportunities to intervene in 1-dSL linked diseases through targeting different signaling arms of the UPR.

## INTRODUCTION

Sphingolipids (SLs) are a class of membrane lipids central to the synthesis of both structural lipids including ceramides, sphingomyelin, and glycosphingolipids and bioactive signaling lipids such as sphingosine-1-phosphate (S1P).(1) In the cell, SLs are synthesized de novo through the condensation of fatty acyl chains to serine, which provides a hydroxyl moiety to which different head groups can be attached. However, during SL synthesis, alanine can be substituted for serine to generate 1-deoxysphingolipids (1-dSLs) that lack the hydroxyl required for lipid functionalization. While the synthesis of 1-dSLs is naturally low, the accumulation of 1-dSLs is cytotoxic(2) and associated with the pathogenesis of numerous retinopathies and neuropathies including type I and type II diabetes, Macular Telangiectasia (MacTel), and hereditary sensory neuropathy type 1 (HSAN1).(3–6)

Gain of function mutations in the first enzyme of the SL biosynthetic pathway, serine palmitoyl transferase (SPT), lead to accumulation of 1-dSL and subsequent neurotoxicity in HSAN1 and retinal degeneration in MacTel.(3, 4) While pathologic SPT mutations are very rare (only a few hundred cases worldwide), MacTel is a more common disease with a prevalence reported up to ~1:1000.(7) MacTel has an extremely heterogenous genetic architecture that converges on a shared metabolic phenotype involving reductions in circulating levels of serine.(3) This reduction in serine drives an elevation of 1-dSLs that correlates with the severity of retinal degeneration. The accumulation of 1-dSLs has also been associated with the onset of peripheral neuropathy and retinopathy in diabetes, where elevated levels are a strong predictive risk factor for type II diabetes(5, 8, 9) and correlate with peripheral neuropathy implicated in type I diabetes.(6)

Intracellular accumulation of 1-dSLs impacts a variety of cellular processes including mitochondrial function(10), lipid body formation(11), protein folding(12), autophagy(13), cytoskeletal reorganization(14), endocytosis(15), calcium handling(16), and ER stress(11, 17). Furthermore, 1-dSLs have been suggested to induce cell death through atypical cell death programs in fibroblast, liver, and neuroblastoma cell lines.(2) 1-dSL toxicity can also vary between the same cells cultured under different conditions, underscoring the sensitivity of 1-dSL toxicity to multiple biological factors.(15) A meta-analysis suggests that neuronal cells are more sensitive to 1-dSLs than other cell types(18), consistent with the pathogenic neuropathy observed in response to 1-dSL accumulation in diseases such as HSAN1 and diabetes. However, despite the clear link between 1-dSLs and cytotoxicity, the pathologic mechanism of 1-dSL toxicity in complex tissues such as the retina remains unclear.

Here, we sought to define a pathologic mechanism for 1-dSL-induced retinopathy using human retinal organoids (ROs). ROs derived from human induced pluripotent stem cells (iPSCs) are functional retinal tissues that recapitulate key aspects of the cytoarchitecture, cellular diversity, and function of the human retina.(19–21) We previously established 1-dSL toxicity in ROs as a model for the degenerative retinal disease MacTel.(3) Late stages of MacTel are characterized by a loss of both photoreceptors and Müller glia in the retinal macula.(22) We found that elevation of 1-dSL concentrations caused apoptosis in photoreceptors and could be rescued through treatment with drugs that directly alter lipid metabolism, including the dyslipidemia drug, fenofibrate.(3) The cytotoxic effect of 1-dSLs is largely attributed to the accumulation of the metabolite 1-deoxydihydroceramide (1-dDHCer)(3, 12), which is synthesized from 1-deoxysphinganine (1-dSA) by Ceramide Synthases (CERSs) in the endoplasmic reticulum (ER)(**Fig. 1A**).(23) Subcellular localization studies using labelled 1-dSA in fibroblasts shows that 1-dSA accumulates in the ER, Golgi, and mitochondria resulting in compromised organelle structure.(10, 24, 25) Further, 1-dSA accumulation induces activation of ER stress-responsive genes including *XBP1s* and the pro-apoptotic transcription factor *DDIT3/CHOP* in other cell types.(10, 26) This suggests that ER stress and impaired ER regulation could contribute to the pathogenic mechanism of 1-dSA-induced retinal toxicity.

**Figure 1.**
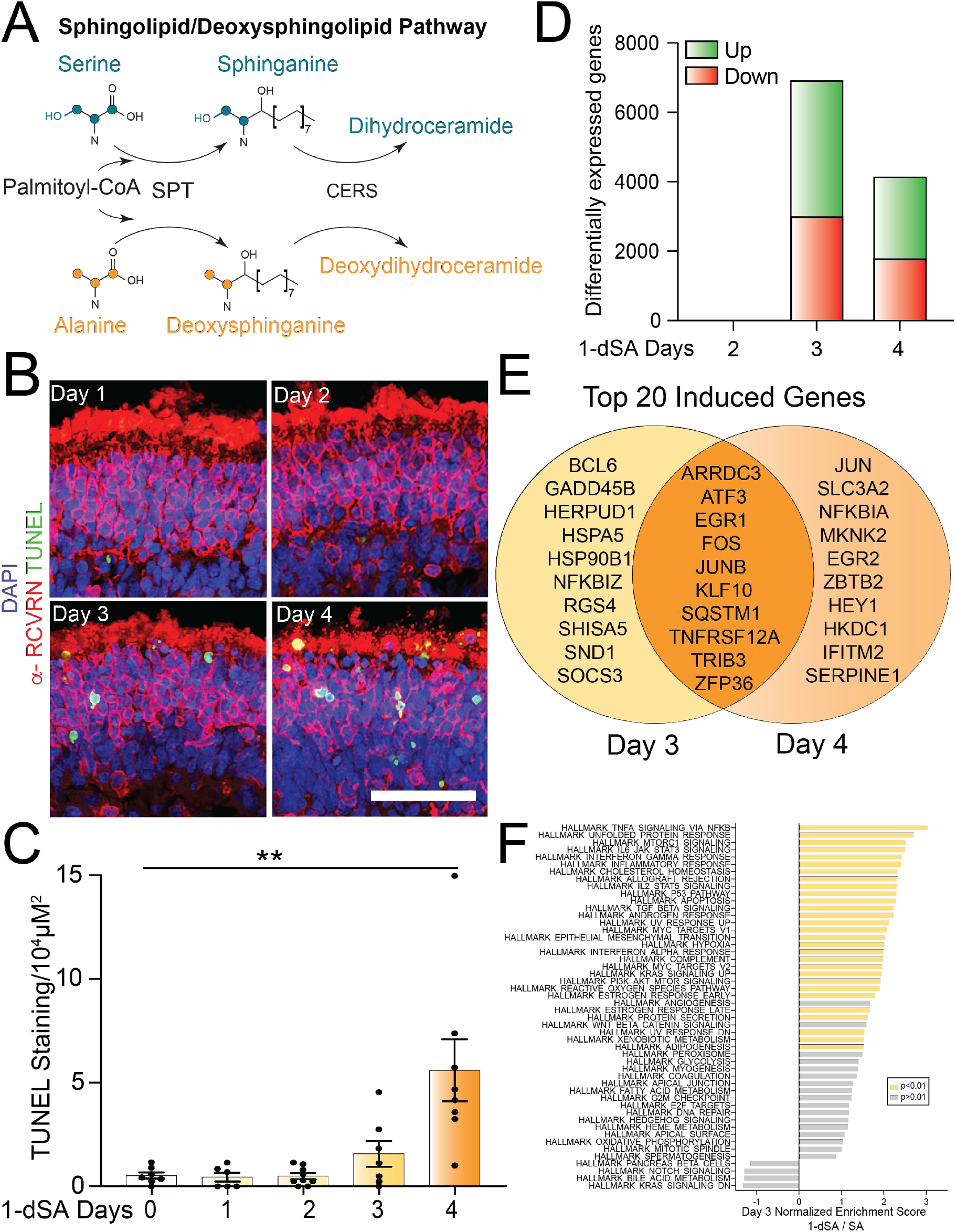
1-dSA-induced transcriptional remodeling precedes cell death. **A**. Schematic of the 1-deoxysphingolipid (1-dSL) biosynthesis pathway. **B**. Immunostaining of the photoreceptor marker α-RCVRN, the cell death marker TUNEL stain, and nuclear DAPI stain in retinal organoids treated with 1-dSA (1 μM) for 1-4 days. Scale bar is 25μm. **C**. Quantification of TUNEL staining from images as shown in (**B**). Error bars show SEM for n=7. **p<0.01 for unpaired Student’s t-test. **D**. Quantification of differentially expressed mRNA transcripts of retinal organoids treated with 1-dSA for 2, 3, and 4 days measured by RNA-seq (n = 5 per condition). **E**. Venn diagram showing the distinct and shared transcripts among the top 20 mRNA transcripts with the highest p values showing increased expression from RNAseq data presented in (**D**). **F**. Enrichment of MSigDB Hallmark pathways in RNAseq data of ROs treated with 1-dSA relative to SA for 3 days (n = 5 per condition). Pathways with enrichment of p-adj < 0.01 are highlighted in yellow.

The unfolded protein response (UPR) is the primary stress-responsive signaling pathway responsible for regulating cellular physiology during ER stress and comprises three signaling pathways activated downstream of the ER stress sensing proteins IRE1, PERK, and ATF6.(27, 28) In response to ER stress, these pathways induce translational and transcriptional signaling that function to both alleviate the ER stress and promote adaptive remodeling of diverse biological pathways involved in cellular functions including protein secretion, lipid synthesis, and calcium regulation.(28, 29) However, chronic UPR activation induces maladaptive signaling, primarily through the PERK-dependent upregulation of pro-apoptotic factors such as *DDIT3/CHOP*, that induce cell death in response to sustained, unresolvable ER stress.(28, 30) Dysregulated UPR signaling is implicated in the onset and pathogenesis of numerous diseases, including many retinal diseases.(31, 32) Deficiencies in ATF6 signaling lead to impaired cone photoreceptor development implicated in the disease achromatopsia.(29, 33) Further, dysregulated signaling through all three UPR pathways is associated with retinal degeneration in diseases such as retinitis pigmentosa.(32–35) This suggests that the retina, and specifically photoreceptor cells within the retina, are highly sensitive to imbalances in UPR signaling.

In this study, we used bulk and single nucleus RNA-seq of mature human ROs to define pathologic mechanisms that contribute to 1-dSL-induced toxicity. We found that 1-dSA treatment induced activation of UPR signaling pathways in photoreceptors and Müller glia, while only minimally impacting gene expression in other cell types. Intriguingly, adaptive IRE1/XBP1s and ATF6 signaling is only observed transiently at early stages of 1-dSA treatment, with activity decreasing at later stages when photoreceptor death is prevalent. In contrast, signaling through the pro-apoptotic PERK arm of the UPR and the related integrated stress response (ISR)(36) is observed in photoreceptors throughout the 1-dSA treatment paradigm. This suggests that deficient signaling through the adaptive IRE1/XBP1s and ATF6 UPR pathways and sustained signaling through the pro-apoptotic PERK/ISR UPR pathway contribute to 1-dSA-induced retinal toxicity. Consistent with this, pharmacologic inhibition of PERK/ISR signaling reduces 1-dSA toxicity in retinal organoids by suppressing the expression of pro-apoptotic and inflammatory genes. In contrast, pharmacologic inhibition of ATF6 activity increases 1-dSA toxicity, while enhancing activation of this UPR signaling pathway mitigates 1-dSA induced photoreceptor cell death. Collectively, our results demonstrate that imbalanced signaling through the UPR, most notably the PERK and ATF6 signaling arms, is a key pathogenic mechanism in 1-dSA-induced retinal toxicity. Further, our results indicate that pharmacologic interventions which correct imbalanced UPR signaling present new opportunities to therapeutically attenuate the retinal degeneration associated with 1-dSL toxicity in diseases such as MacTel or diabetic retinopathy.

## RESULTS

### Treatment with 1-dSA induces time-dependent toxicity in retinal organoids

We sought to define the pathologic mechanisms that contribute to 1-dSA-induced toxicity in ROs. We initially monitored apoptosis in ROs treated with 1-dSA (18:0) for 1-4 days using TUNEL staining. No TUNEL staining was observed following 1-2 days of treatment, with a modest increase following 3 days of treatment (**Fig. 1B,C**). Retinal cell death was prominently observed at day 4. The majority of TUNEL staining was observed in the outer nuclear layer (ONL), which primarily consists of photoreceptors (**Fig. S1A**). This is consistent with the previously observed sensitivity of photoreceptors to 1-dSA. (3, 37)

To define pathways involved in the initial steps of 1-dSA toxicity, we performed whole transcriptome RNA-seq on ROs treated with 1-dSA for 2, 3, or 4 days. We observed only 1 differentially expressed gene (DEG) following 2 days of treatment, with substantially higher DEGs observed following 3 or 4 days of treatment (**Fig. 1D**, **Tables S1-3**). The highest number of DEGs were observed following 3 days of treatment, corresponding to the modest increase in toxicity observed at this timepoint (**Fig. 1B,C**). The transcriptional response observed upon treatment of ROs with 1-dSA for 3 days was distinct from that observed in ROs treated with a non-toxic SL, sphinganine (**Fig. S1B,C**, **Table S2**), indicating that the observed transcriptional response was specific to deoxy-derived SLs rather than a broader response to exogenous lipids. DEGs observed in ROs treated for 3 or 4 days with 1-dSA showed significant overlap, although many genes were also found to be differentially expressed between these two time points (**Fig. 1E, Fig. S1D**). Gene set enrichment analysis (GSEA) demonstrated enrichment of pathologic pathways involved in inflammation, apoptosis, and UPR signaling at both 3 and 4 days of treatment (**Fig. 1F**, **Fig. S1E**). This suggests an important role for these biological pathways in 1-dSA retinal toxicity.

### Photoreceptors and Müller glia are selectively sensitive to 1-dSA in ROs

To gain further insights into the cell type-specific mechanisms of 1-dSA-induced toxicity, we performed single nucleus RNAseq (snRNAseq) on iPSC-derived ROs treated with or without 1-dSA for 3 days – a timepoint when substantial transcriptional remodeling is first observed (**Fig. 1B-D**). Integrated clustering of control and 1-dSA-treated ROs showed overlapping groups of mature retinal cell types including rod and cone photoreceptors, Müller glia, accessory cell types, and immature and progenitor cells (**Fig. 2A**, **Fig. S2A**). When we compared the size of the transcriptional response in each cell type, we observed that 1-dSA substantially increased DEGs in Müller glia and cone and rod photoreceptors, with minimal effects on other retinal cell types (**Fig. 2B, Table S4)**. GSEA analysis of Müller glia and cone/rod photoreceptors showed that all three of these cell types demonstrated enrichment for pathways involved in inflammation, apoptosis, and UPR signaling (**Fig. S2B-D**). This is consistent with the increase in these pathways observed by bulk RNAseq of 1-dSA-treated ROs (**Fig. 1F**).

**Figure 2.**
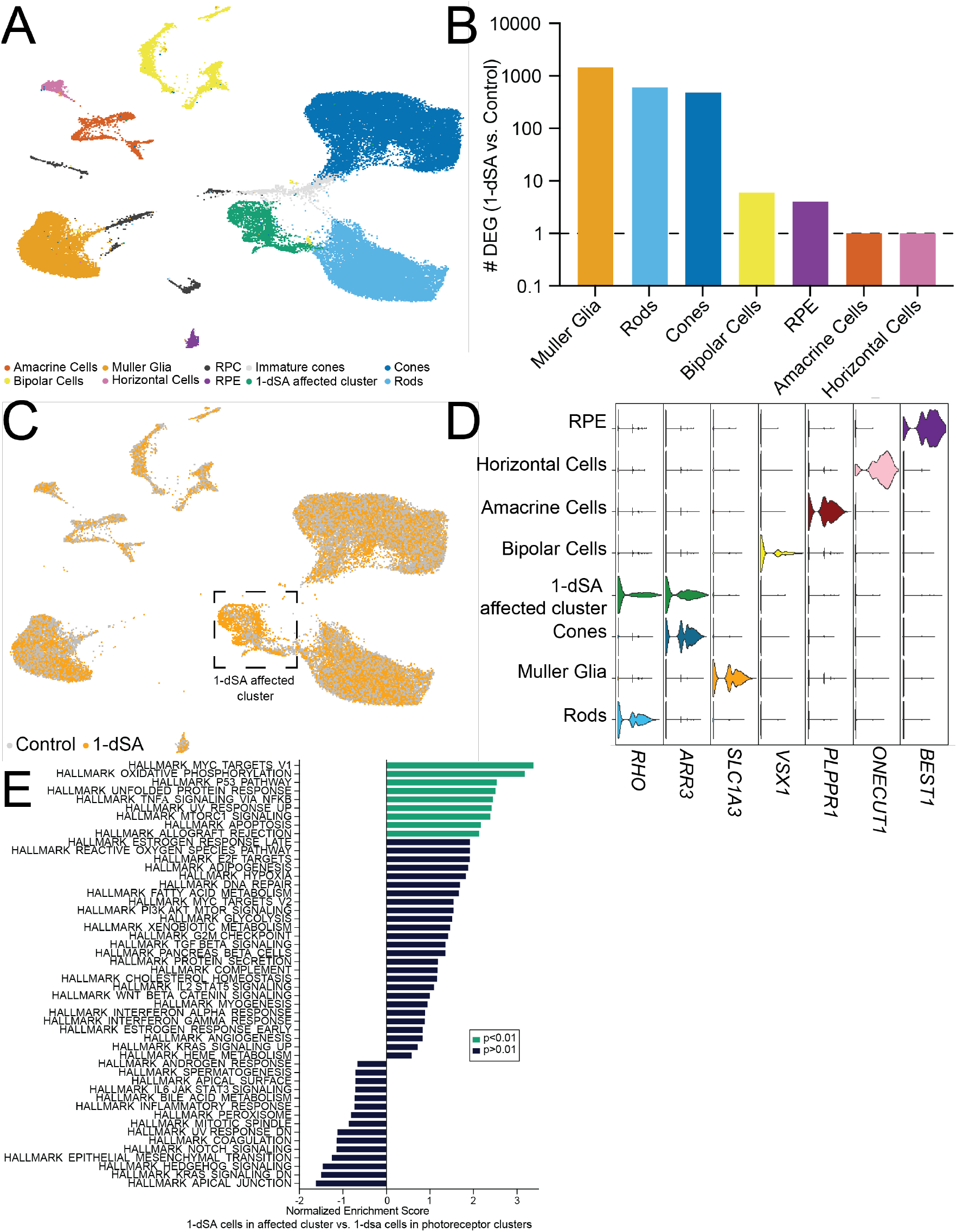
Photoreceptors and Müller glia show a transcriptional response to 1-dSA. **A**. UMAP of unbiased clustering of snRNAseq data from ROs treated with control or 1-dSA (1 μM) for 3 days colored by cluster (n=2 organoids per condition were used for snRNAseq). Clusters were assigned a retinal cell type based on previously defined markers. **B**. Bar graph showing the total number of differentially expressed genes (DEGs) between control and 1-dSA (1 μM; 3 days) organoids for each mature retinal cell cluster identified in (**A**). **C**. UMAP of all cells in snRNAseq data colored by treatment. 1-dSA-treated organoids are in orange; control-treated organoids are in grey. **D**. Violin plot of mature retinal cell type marker expression by clusters, as defined in in (**A**). **E**. Enrichment plot of differentially expressed pathways between 1-dSA treated cells in the affected cluster compared to 1-dSA treated cells in the combined rod and cone clusters. Pathways enriched in the 1-dSA affected cluster (p<0.01) are highlighted in green.

When we compared cell type population sizes between 1-dSA and control conditions, we observed that 1-dSA increased a specific population of cells, herein referred to as the 1-dSA affected cluster (**Fig. 2C**), which clustered closely to photoreceptors. Cells in this cluster expressed markers of cones (e.g., *ARR3*) and rods (e.g., *RHO*), but not markers of other retinal cell types (**Fig. 2D, Fig. S2E**). The increase in cells observed in the 1-dSA affected cluster also corresponded with reductions in cone photoreceptors (**Fig. S2F**), further indicating that this population reflects an altered state of photoreceptor cells. A comparison of gene expression between the 1-dSA affected cluster, and the rod and cone clusters within the 1-dSA treatment shows further enrichment of multiple stress-responsive pathways including oxidative phosphorylation, P53-mediated apoptosis, and the UPR (**Fig. 2E**). This suggests that cells within the 1-dSA affected cluster are likely photoreceptors primed for death. This is consistent with TUNEL staining showing the majority of cell death occurring in the photoreceptors of the RO ONL (**Fig. S1A**).

### The three arms of the UPR are differentially activated in retinal cell types

GSEA of our bulk RNAseq identified the unfolded protein response (UPR) as a prominent stress pathway activated in retinal organoids treated with 1-dSA for both 3 (**Fig. 1F**) or 4 days (**Fig. S1E**). We similarly observed significant upregulation of the UPR in photoreceptors and Müller glia (**Fig. S3A,B**), the cell types most impacted by 1-dSA treatment. Based on the prominent and timely activation of the UPR in 1-dSA treated ROs, and previous studies that have localized the conversion of 1-dSA to the toxic species 1-dDHCer by CERS in the ER we hypothesized that the ER stress responsive UPR plays an important role in 1-dSA mediated toxicity.

The UPR comprises three signaling pathways activated downstream of the ER stress sensing proteins PERK, IRE1, and ATF6 (**Fig. 3A**).(28) In response to ER stress, these pathways promote transcriptional remodeling through the activation of the UPR-associated transcription factors ATF4, XBP1s, and cleaved ATF6, respectively.(27, 28) We used sets of genes differentially regulated downstream of these three UPR signaling pathways(38) to define the relative activity of PERK/ATF4, IRE1/XBP1s, and ATF6 signaling in ROs treated with 1-dSA for 3 days. Bulk RNA-seq showed that 1-dSA induced expression of target genes associated with all three UPR arms, indicating that all three UPR signaling pathways are activated at this timepoint (**Fig. 3B, Fig. S3C**). Further, snRNAseq showed increased expression of target genes regulated by PERK/ATF4 (e.g., *DDIT3, ATF4*) and ATF6 (e.g., *CALR, HSPA5, HERPUD1*) in the 1-dSA affected cluster following three days of treatment (**Fig. 3C,D**). The canonical XBP1s target *DNAJB9* showed the highest levels of expression in the 1-dSA-affected cluster (**Fig. S3D**); however, other XBP1s target genes were overall poorly detected and could not be used to define patterns of IRE1 activity in individual retinal cell types (**Fig. S3E**). This suggests that all three UPR arms are activated in the 1-dSA affected cluster under these conditions. In contrast, Müller glia showed basally higher expression of some ATF6 target genes (e.g., *CALR, HSPA5*) with 1-dSA-induced increases in the expression of other ATF6 targets such as *PDIA6* (**Fig. 3D**). PERK/ATF4 target genes were not induced in these cells (**Fig. 3C**). No other retinal cell type showed significant activation of any UPR signaling pathways in our snRNAseq data. These results indicate that distinct UPR signaling was preferentially induced in the 1-dSA affected cluster of photoreceptors and Müller glia in ROs treated with 1-dSA for 3 days.

**Figure 3.**
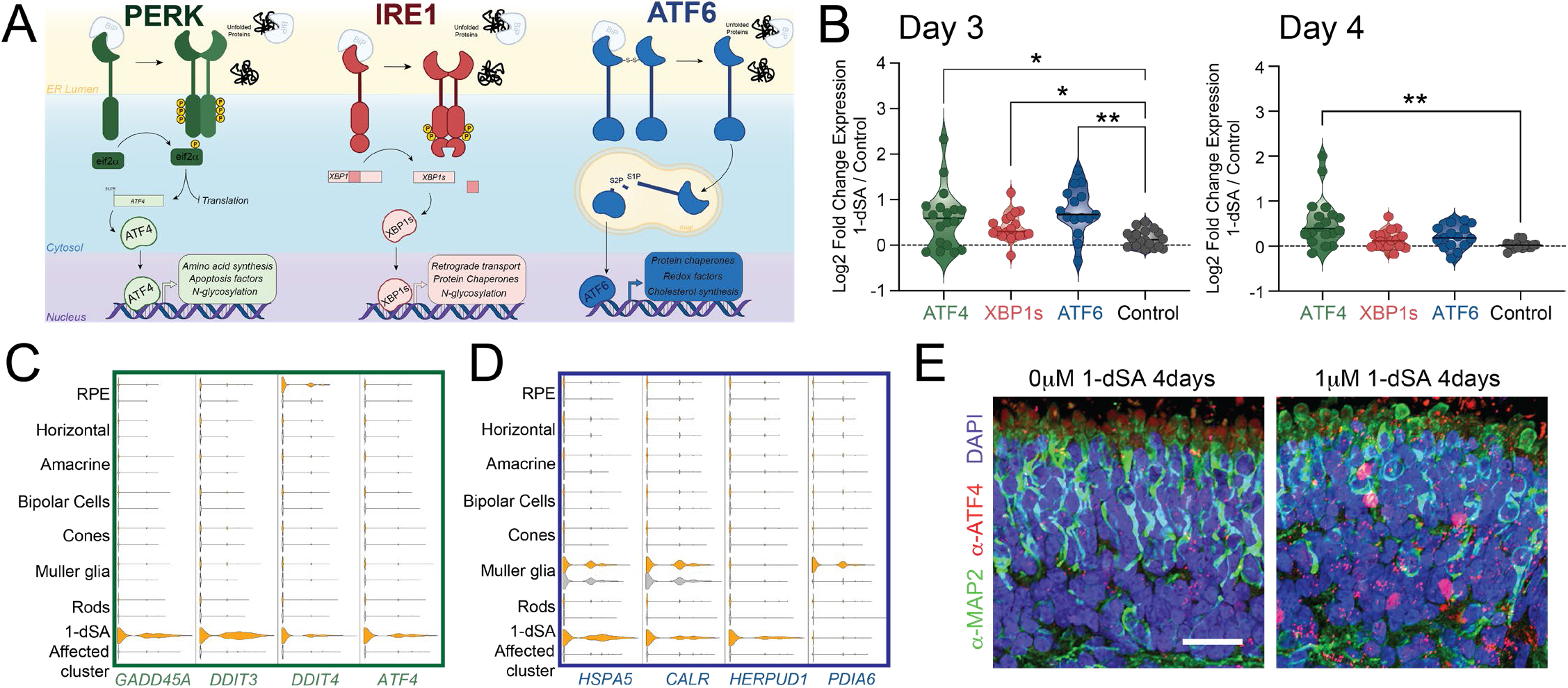
1-dSA differentially activates Unfolded Protein Response signaling pathways in specific retinal cell types. **A**. Illustration showing the transcriptional and translational remodeling which occurs downstream of PERK, IRE1, and ATF6 activation. **B**. Quantification of RNAseq fold changes of gene targets of the UPR-induced transcription factors ATF4, XBP1s, and ATF6 in ROs treated with 1-dSA (1 μM) relative to control for 3 or 4 days. *p<0.05, **p<0.01 for Brown-Forsythe and Welch ANOVA tests compared to control gene set. (n=5, per condition for each dataset). **C**. Violin plot of gene targets of ATF4 in snRNAseq dataset (n=2 organoids per condition were used for snRNAseq) separated by cluster and treatment. 1-dSA-treated organoids (3 days; 1 μM) are in orange; control organoids are in grey. **D**. Violin plot of gene targets of ATF6 in snRNAseq dataset separated by cluster and treatment. 1-dSA treated organoids (3 days; 1 μM) are in orange; control organoids are in grey. **E**. Immunostaining of the photoreceptor marker α-MAP2 (green), α-ATF4 (red), and nuclear stain DAPI (blue) in ROs treated with 1-dSA (1 μM) for 4 days.

Intriguingly, while ROs treated for 3 days with 1-dSA showed activation of genes regulated by PERK/ATF4, IRE1/XBP1s, and ATF6, RNAseq data from ROs treated for 4 days with 1-dSA showed preferential activation of the PERK/ATF4 signaling arm of the UPR (**Fig. 3B**). Despite seeing increased expression of select IRE1/XBP1s and ATF6 target genes, the majority of targets associated with these pathways were not induced at day 4. This suggests that the IRE1/XBP1s and ATF6 pathways are not robustly activated at this timepoint. We confirmed prominent increases of ATF4 protein expression following four days of 1-dSA treatment by immunostaining (**Fig. 3E**). Collectively, our results indicate that all three arms of the UPR are activated at day 3 of 1-dSA treatment, a timepoint before significant RO toxicity is observed (**Fig. 1B,C**). However, only PERK/ATF4 signaling persists at day 4 when higher levels of TUNEL staining is observed.

### PERK signaling promotes 1-dSA toxicity in ROs

Chronic or hyperactive PERK/ATF4 signaling induces apoptosis in multiple models through increased expression of pro-apoptotic factors such as *DDIT3/CHOP*.(30, 39, 40) Our observation that PERK/ATF4 target genes show persistent activation in 1-dSA treated ROs suggests that hyperactive PERK signaling could contribute to the observed toxicity in this model. We monitored toxicity in ROs treated with 1-dSA in the presence or absence of two mechanistically distinct PERK signaling inhibitors; GSK2656157 and ISRIB. GSK2656157 is a PERK kinase inhibitor that blocks PERK autophosphorylation required for the activation of this UPR signaling pathway (**Fig. 4A**).(41) Alternatively, ISRIB inhibits PERK signaling downstream of eIF2α phosphorylation.(42–44) Thus, unlike GSK2656157, ISRIB can also inhibit eIF2α phosphorylation and downstream signaling induced by other eIF2α kinases (e.g., GCN2, HRI, PKR) comprising the integrated stress response (ISR).(36, 44) Co-treatment with either inhibitor reduced TUNEL staining in ROs treated with 1-dSA, indicating that PERK signaling contributes to 1-dSA-induced toxicity in this model (**Fig. 4B,C**). ISRIB showed a stronger reduction in toxicity, as compared to GSK2656157, potentially indicating a role for other ISR kinases in this toxicity. Both compounds inhibited ATF4 target gene expression in 1-dSA-treated ROs, confirming their activity (**Fig. S4A, Fig. 4D**). These results indicate that PERK/ISR signaling is an important contributor to photoreceptor death in 1-dSA-treated ROs.

**Figure 4.**
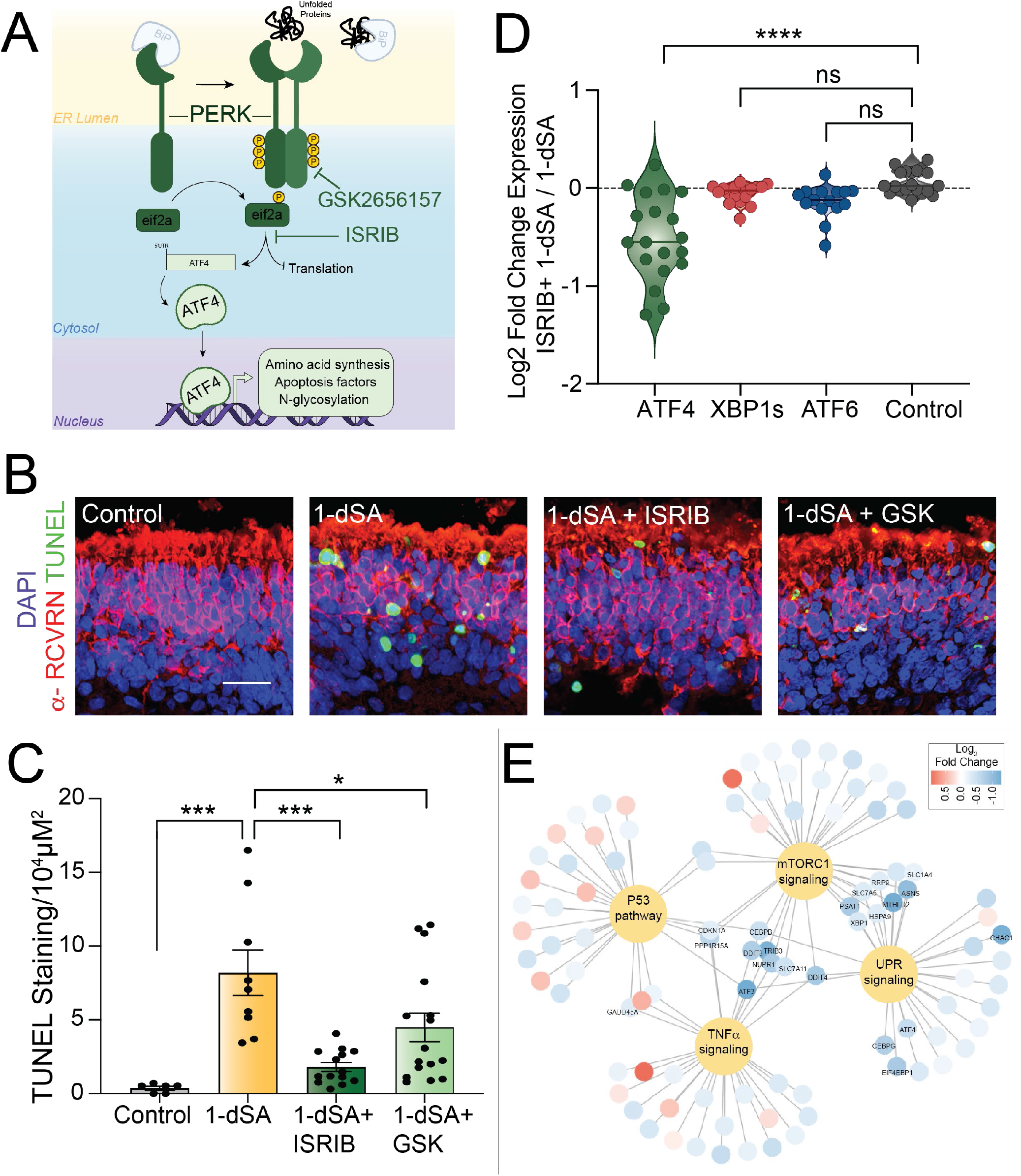
PERK/ISR signaling drives 1-dSA induced cell death. **A**. Illustration showing PERK signaling induced by ER stress. GSK2656157 inhibits PERK autophosphorylation and ISRIB inhibits signaling downstream of eIF2α phosphorylation. **B,C**. Representative images and quantification of TUNEL staining of organoids treated for 4 days with 1-dSA (1 μM) in the presence or absence of the PERK kinase inhibitor GSK2656157 (500 nM) or the PERK signaling inhibitor ISRIB (200 nM). **B**. Scale bar is 25μm. **C**. Error bars show SEM for n=9-15. *p<0.05, ***p<0.005 for ordinary one-way ANOVA. **D**. Quantification of RNAseq fold changes of gene targets of the UPR-induced transcription factors ATF4, XBP1s, and ATF6 in ROs treated with 1-dSA (1 μM) in the presence or absence of ISRIB (200 nM) for 4 days. *p<0.05, **p<0.01 for Brown-Forsythe and Welch ANOVA tests compared to control gene set (n = 5 per condition). **E**. Network graph showing shared transcripts and associated fold changes of Hallmark gene set pathways significantly altered by treatment with 1-dSA.

To further define the involvement of PERK signaling in this process, we performed bulk RNAseq on ROs treated with 1-dSA and/or ISRIB for 4 days. Co-treatment of 1-dSA with ISRIB reduced expression of PERK/ATF4 target genes in this model but did not significantly influence expression of genes regulated by other arms of the UPR (**Fig. 4D**). GSEA shows that ISRIB co-treatment also reduced expression of genes involved in P53-mediated apoptosis, TNFα mediated inflammation, and mTORC1 activation (**Fig. S4B**) – three pathways upregulated by 1-dSA in Müller glia and photoreceptor clusters (**Fig. 2E, Fig. S2B-D**). However, these reductions can be largely attributed to the reduced expression of ATF4-regulated genes including *DDIT3, ATF3*, and *PPP1R15A* (**Fig. 4E, Fig. S4C**), indicating that the downregulation of these other pathways observed by GSEA is likely driven by the suppression of PERK/ISR signaling. Collectively, these results indicate that pharmacologic inhibition of PERK/ISR signaling selectively impacts the transcriptional response to 1-dSA in ROs predominantly by suppressing ATF4 activity.

The above results suggest pharmacologic inhibition of PERK/ISR signaling as a potential strategy to mitigate 1-dSA-induced toxicity. However, PERK/ISR signaling is also implicated in the regulation of amino acids, including serine.(45, 46) Notably, PERK/ISR activation induces the ATF4-mediated expression of key serine biosynthesis enzymes including *PSAT1* and *PHGDH*. Decreased activity of these enzymes reduces serine availability and increases cellular production of 1-dSA and toxic dSLs.(3, 11, 47) This suggests that pharmacologic inhibition of PERK/ISR signaling could exacerbate 1-dSA production by suppressing serine biosynthesis. Consistent with this, ISRIB reduced expression of serine biosynthesis genes in ROs both in the absence and presence of 1-dSA (**Fig. S4D,E**). Thus, while PERK/ISR inhibition blocks 1-dSA toxicity in ROs, it may exacerbate production of toxic 1-dSLs and worsen cellular and tissue damage in the context of human disease.

### ATF6 activity is protective in 1-dSA-treated ROs

The IRE1 and ATF6 signaling arm of the UPR are generally associated with adaptive remodeling of cellular physiology in response to ER stress. This is primarily mediated through the activation of the transcription factors ATF6 (a cleaved product of full-length ATF6) and XBP1s (downstream of IRE1) (**Fig. 5A** and **Fig. S5A**, respectively). These transcription factors induce expression of genes involved in numerous adaptive biological pathways including ER proteostasis maintenance, cellular metabolism, and redox regulation.(29, 48) Our transcriptional profiling showed transient increases in the expression of ATF6, and to a lesser extent IRE1/XBP1s, target genes in ROs treated with 1-dSA for 3 days (**Fig. 3B**). Thus, we sought to define the specific contributions of these UPR signaling pathways in 1-dSA toxicity using pharmacologic inhibitors and activators of these adaptive UPR signaling pathways.

**Figure 5.**
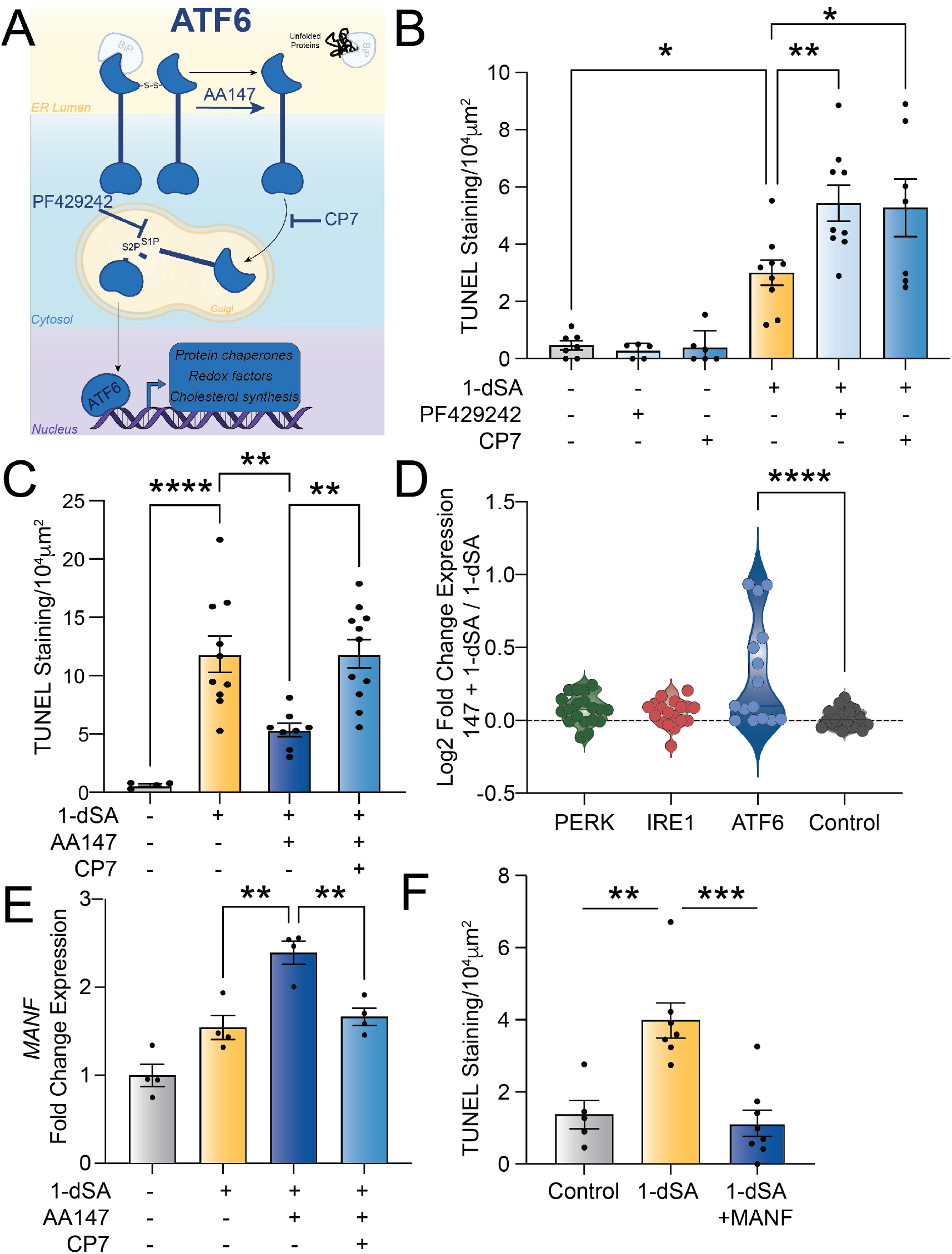
ATF6 promotes retinal cell protection in the context of 1-dSA toxicity. **A**. Illustration of the ATF6 UPR signaling pathway and pharmacologic modulators of this pathway. **B**. Quantification of TUNEL staining of ROs treated with 1-dSA (1μM) for 4 days in the presence or absence of the ATF6 inhibiting compounds Ceapin-A7 (CP7; 7 μM) or PF429242 (10. **C**. Quantification of TUNEL staining of ROs treated with 1-dSA (1 μM) as well as AA147 (10μM) in the presence or absence of Ceapin-7 (CP7). **D**. Quantification of RNAseq fold changes of gene targets of the UPR-induced transcription factors ATF4, XBP1s, and ATF6 in ROs treated with 1-dSA in the presence or absence of ISRIB (200 nM) for 3 and 4 days. ****p<0.0001 for Brown-Forsythe and Welch ANOVA tests compared to control gene set. **E**. qPCR of the ATF6 gene target *MANF \*\**p<0.01 for ordinary one-way ANOVA.

IRE1 was inhibited using compounds 4μ8c and STF-083010, both compounds that directly inhibit IRE1 RNAse activity required for IRE1-dependent XBP1s activation (**Fig. S5A**).(49, 50) We confirmed that co-treatment with these compounds inhibited 1-dSA-dependent increases in *XBP1s* (**Fig. S5B**). 1-dSA co-treatment with the IRE1 inhibitor STF-083010 increased toxicity in 1-dSA-treated cells; however, we did not observe similar increases upon co-treatment with 4μ8c (**Fig. S5C**). Further, pharmacologic activation of IRE1/XBP1s signaling using the IRE1/XBP1s activator compound IXA4(51) upregulated XBP1s levels (**Fig. S5D**) but did not reduce toxicity in 1-dSA-treated retinal organoids (**Fig. S5E**). Collectively, these results suggest that IRE1/XBP1s activity is not prominently involved in influencing 1-dSA toxicity in ROs.

Next, we used pharmacologic activators and inhibitors of ATF6 to probe the importance of this pathway in 1-dSA retinal toxicity. ATF6 activity was inhibited using two compounds, Ceapin-A7 and PF-429242, that block ATF6 activity through two distinct mechanisms (**Fig. 5A**). Ceapin-A7 inhibits ATF6 activation by preventing its trafficking to the Golgi for proteolytic activation. (52) In contrast, PF-429242 inhibits S1P proteolytic activity required for release of the active, cleaved ATF6 transcription factor.(53) We confirmed that co-treatment with either Ceapin-A7 or PF-429242 inhibited 1-dSA-dependent induction of the ATF6 target gene *HSPA5/BiP*, confirming the activity of these compounds in ROs (**Fig. S5F**). We also observed that co-treatment with either of these two ATF6 inhibitor increased toxicity in 1-dSA-treated ROs (**Fig. 5B**, **Fig. S5G**), suggesting that 1-dSA-induced ATF6 activation protects against cell death.

To further probe the contributions of ATF6 activation in 1-dSA-induced toxicity, we employed AA147 – a pharmacologic activator of ATF6 transcriptional activity (**Fig. 5A**).(54) AA147 promotes ATF6 activation through a mechanism involving increased trafficking to the Golgi for proteolytic activation.(55) We confirmed that co-treatment with AA147 increased expression of the ATF6-target gene *HSPA5/BiP* in the presence of 1-dSA (**Fig. S5H**). Co-treatment with the ATF6 inhibitor Ceapin-A7 blocked AA147-dependent increases in *HSPA5/BiP* expression, confirming this compound increased *BiP* expression through an ATF6-dependent mechanism. AA147 treatment increased survival of ROs challenged with 1-dSA (**Fig. 5C**, **Fig. S5I**). This protection was lost upon co-treatment with Ceapin-A7, indicating that AA147 increased protection through an ATF6-dependent mechanism.

Bulk RNA-seq profiling of ROs treated with AA147 and 1-dSA showed increased expression of multiple ATF6 target genes (**Fig. 5D**). IRE1/XBP1s target genes were not induced by AA147, reflecting the selectivity of this compound for ATF6 activation (**Fig. 5D**).(54) PERK target genes were also not significantly altered in AA147-treated cells co-treated with 1-dSA, indicating that AA147-dependent ATF6 activation did not reduce 1-dSA toxicity by suppressing PERK/ISR signaling.

ATF6 regulates the expression of multiple genes known to be protective in the retina, including the secreted neurotrophic factor MANF – a protein primarily expressed in Müller glia (**Fig. S5J**).(56) Extracellular MANF has previously been shown to protect the retina from diverse types of insults.(55–59) AA147 significantly increases expression of MANF in 1-dSA-treated ROs (**Fig. 5E**). This is suppressed by co-treatment with Ceapin-A7, confirming that AA147 increases *MANF* expression through an ATF6-dependent mechanism. This indicates that ATF6-dependent increases in MANF could contribute to the protection observed upon treatment with AA147. Consistent with this, administration of recombinant MANF reduces 1-dSA-induced toxicity in ROs (**Fig. 5F**, **Fig. S5K**). These results suggest that AA147-dependent ATF6 activation protects against 1-dSA-induced toxicity through the upregulation of adaptive, protective target genes such as MANF.

## DISCUSSION

The accumulation of cytotoxic 1-dSLs is linked to multiple retinopathies and neuropathies(3–6), however the mechanisms by which 1-dSLs impact retinal cell function are poorly understood. Here, using transcriptomic profiling of ROs treated with 1-dSA across time and at a single cell resolution, we characterized a broad cellular response with a pronounced activation of the ER stress responsive UPR. We subsequently validated the functional role of the UPR in 1-dSL toxicity by utilizing compounds that selectively inhibit or activate individual UPR signaling arms. This approach showed that PERK/ISR signaling mediates cell death through activation of pro-apoptotic and inflammatory pathways, whereas ATF6 promotes cell survival through mechanisms including upregulation of the secreted neurotrophic factor MANF. Although the activation of the pro-apoptotic PERK arm of the UPR is maintained throughout the 1-dSA treatment paradigm, we observed that activation of the pro-survival ATF6 arm is activated primarily at early stages prior to elevated cell death. This indicates that imbalanced signaling through these pro-apoptotic and adaptive signaling arms of the UPR is an important contributor to 1-dSL-induced retinal toxicity.

Our results suggest that enhancing/prolonging ATF6 signaling offers a unique opportunity to mitigate pathologic photoreceptor death induced by 1-dSA treatment. Pharmacologic inhibition of ATF6 exacerbates 1-dSL toxicity, demonstrating an adaptive, protective role for ATF6 signaling in the context of 1-dSL toxicity. Pharmacologically enhancing ATF6 activity with AA147 attenuated 1-dSL toxicity, indicating that increasing and maintaining signaling through this adaptive pathway could mitigate the toxicity associated with disease. ATF6 signaling is well established to be important for regulating retinal development and health. Hypomorphic mutations in *ATF6* impair cone photoreceptor differentiation in the disease achromatopsia.(32) As we observed, pharmacologic ATF6 activation can rescue this deficiency and restore cone photoreceptors in iPSC models of this disease.(60) Alternatively, deficiencies in ATF6 activation induced by environmental insults or aging contribute to retinal degeneration associated with other diseases including retinitis pigmentosa and cone-rod dystrophy.(34, 61) The protection afforded by ATF6 in the retina is likely mediated through its regulation of multiple adaptive genes including the ER chaperones BiP and the neurotrophic factor MANF.(29, 62, 63) Each of these ATF6-regulated genes have been shown to protect photoreceptors and/or Müller glia against diverse types of insults including ER-stress induced apoptosis(34, 64, 65), oxidative stress(66), and age-related retinal inflammation.(67) We show that the exogenous addition of MANF alone is sufficient to rescue cell death in 1-dSL toxicity, indicating a likely role for ATF6-dependent regulation of *MANF* expression in the protection observed for AA147 and providing an additional prospective treatment to mitigate 1-dSL-associated retinal toxicity. However, it is important to note that AA147-dependent ATF6 activation likely mediates its protection through the regulation of multiple adaptive factors, underscoring the unique potential for pharmacologically targeting this UPR signaling pathway to mitigate 1-dSL toxicity in complex tissues such as the retina.

Pharmacologic inhibition of PERK kinase activity with GSK2656157 reduced 1-dSL toxicity, whereas inhibition of eIF2α signaling downstream of PERK using the compound ISRIB provided more substantial protection. This suggests that other eIF2α kinases of the ISR, apart from PERK, may also be involved in 1-dSA mediated toxicity. Consistent with this, serine deprivation and exogenous 1-dSA addition can activate ISR kinases such as GCN2 and PKR that promote eIF2α phosphorylation and ATF4 transcriptional activity(68, 69). As ATF4 regulates expression of serine synthesis genes, 1-dSL-induced activation of PERK and other eIF2α kinases may function as an adaptive response for regulating intracellular serine levels. However, since PERK/ISR signaling is important for regulating serine synthesis, pharmacologic targeting of these pathways for 1-dSL-associated disorders such as MacTel is unlikely to be a promising path forward for treating patients, as downregulation of serine synthesis genes would exacerbate the underlying pathology of elevated 1-dSLs.

Using snRNAseq, we show that within retinal tissue there are distinct cell specific responses to 1-dSL, with photoreceptors and Müller glia demonstrating the most pronounced response. This is consistent with the pathogenesis of MacTel where these cellular subtypes are also the retinal cell types most profoundly impacted in this disease.(22) It remains unclear why photoreceptors and Müller glia are uniquely reactive to 1-dSLs. One possibility is the elevated conversion of 1-dSA to the toxic 1-dhCER in photoreceptors and Müller glia through elevated expression of CERS family enzymes. However, we do not observe elevated expression of total CERS in photoreceptors and Müller glia, nor do we observe cell-specific expression of CERS family isoforms that correspond to cell-specific toxicity. Another hypothesis is that cell-specific toxicity is dependent on the unique physiological demands of each cell type. If so, future work determining the physiological consequences of elevated 1-dSLs will be essential to understand the genesis of organelle dysfunction. Regardless, our analysis here suggests that the disruption of the ER in both photoreceptors and Müller glia is a prime event in 1-dSL toxicity in the retina.

Collectively, our results indicate that imbalanced signaling through the pro-apoptotic PERK/ISR and adaptive ATF6 arms of the UPR in two key retinal cell types, Müller glia and photoreceptors, is a contributing factor in dictating 1-dSA toxicity in ROs. This provides a framework to better understand how 1-dSA promotes toxicity of the retina and peripheral neurons in diverse diseases including MacTel, diabetes, and HSAN1. Further, our results identify pharmacologic enhancement of ATF6 activity and/or exogenous addition of the ATF6-regulated neurotrophic factor MANF as potential strategies to promote adaptive remodeling of the retina to mitigate pathology associated with increases of 1-dSLs.

## METHODS

### Organoid Generation and Maintenance

The human induced pluripotent stem cell (hiPSC) line used was derived from peripheral blood mononuclear cells from a female. Reprogramming was performed by the Harvard iPS core facility using sendai virus for reprogramming factor delivery. All cell lines were obtained with verified normal karyotype and contamination-free. hiPSC were maintained on Matrigel (BD Biosciences) coated plates with mTeSR+ medium (STEMCELL Technologies). Cells were passaged every 3-4 days at approximately 80% confluence. Colonies containing clearly visible differentiated cells were marked and mechanically removed before passaging.

Retinal organoids were differentiated from hiPSCs between passage 10 and 20. Retinal organoids were initiated and differentiated as previously described.(70) Following week 4 of differentiation, organoids were cultured in rotating cell suspension until week 18. Mature retinal organoids were cultured in Retinal Differentiation Media plus 10% FBS, 100 μM Taurine, and 2 mM Glutamax starting at week 8 of differentiation. Fully mature retinal organoids were assayed between 26 and 30 weeks post differentiation.

### Cell culture treatments

Lyophilized lipids, 1-deoxy-sphinganine (Avanti; cat 860493) and sphinganine (Avanti; cat 860498), were resuspended to stock concentrations at 5 mM in EtOH and subsequently added to retinal organoid culture media at a concentration of 1μM. For control conditions equivalent amounts of EtOH were added to media. ISRIB (Sigma; cat SML0843) was administered at 200nM. GSK2656157 (Bio Vision; cat 9466) was administered at 500 nM. AA147 was obtained from the Kelly Lab at Scripps Research and resuspended in dimethyl sulfoxide (DMSO); organoids were administered at 10 μM daily. PF429242 (Sigma-Aldrich; cat SML0667) was resuspended in water and administered at 10 μM. CP7 was obtained from the Walter Lab at UCSF, resuspended in DMSO, and administered at 7 μM. 4μ8C (EMD millipore; cat 412512) and STF-083010 (Sigma; cat 412510) were resuspended in DMSO and administered at 32 μM. For drug experiments 1-dSA and drugs were added to organoid culture media at concurrent times. Organoids were cultured in experimental conditions for 4 days unless otherwise stated with a condition specific media change at day 2. For GSK2656157 and ISRIB experiments, drugs were added at day 0 and day 2. For all other drug experiments, drugs were added daily.

### Immunohistochemistry and TUNEL staining

Organoid tissue fixed in 4% PFA in PBS for 10 mins, washed in PBS, and then in 20% sucrose in PBS overnight. Tissues were embedded in O.C.T compound and frozen. Cryosectioning was done at 12 μm and slices were mounted on glass polylysine coated slides. Prior to primary antibodies samples were blocked with 5% donkey serum in PBS. Primary antibodies were added to samples at 4°C and incubated overnight. Following primary antibodies samples were washed 3 × 10 mins in PBS. Secondary antibodies were added at room temp for 2 hours. Dapi was added at 1:1000 in PBS for 10 mins following secondary antibodies.

Primary antibodies: rabbit anti Recoverin (1:500, Millipore AB5585), mouse anti Map2 (1:500, BD Bioscience 556320), ATF4 (1:200 Cell Signaling 11815) (Secondary antibodies: donkey anti rabbit alexafluor 555 (1:1000, Invitrogen 31572), donkey anti rabbit alexafluor 488 (1:1000, Invitrogen 21206), donkey anti mouse alexafluor 555 (1:1000, Invitrogen 31570), donkey anti mouse alexafluor 488 (1:1000, Invitrogen 21202). TUNEL staining was performed using In Situ Cell Death Detection Fluorecein kit (Sigma cat# 11684795910) prior to addition of anti-recoverin primary antibody. Overlapping TUNEL positive staining and DAPI staining within a recoverin positive cell was counted as cell death within a photoreceptor. Cell death was normalized to the area of recoverin staining in the retinal organoid.

### RNA isolation and quantitative RT-PCR

Total RNA was purified from frozen tissues using Trizol Reagent (Life Technologies) according to the manufacturer’s instructions. First-strand cDNA was synthesized from 400ng of total RNA using High-Capacity Reverse Transcriptase kit (Applied Biosystems) according to the manufacturer’s instructions. Individual 10 μl SYBR Green Master Mix (Applied Biosystems) real-time PCR reactions consisted of 2 μl of diluted cDNA, 5 μl of Power Up SYBR Green (Applied Biosystems), and 1 μl of each 5 μM forward and reverse primers. The PCR was carried out on 384-well plates on a Quant Studio Real Time PCR system (Applied Biosystems) using a three-stage program: 95 °C for 10 min, 40 cycles of 95 °C for 20 s, 60 °C for 20 s and 72 C for 20 s. PCR data for intergene comparison were corrected for primer efficiency. Samples were normalized to internal loading control, 36B4.

#### Primers

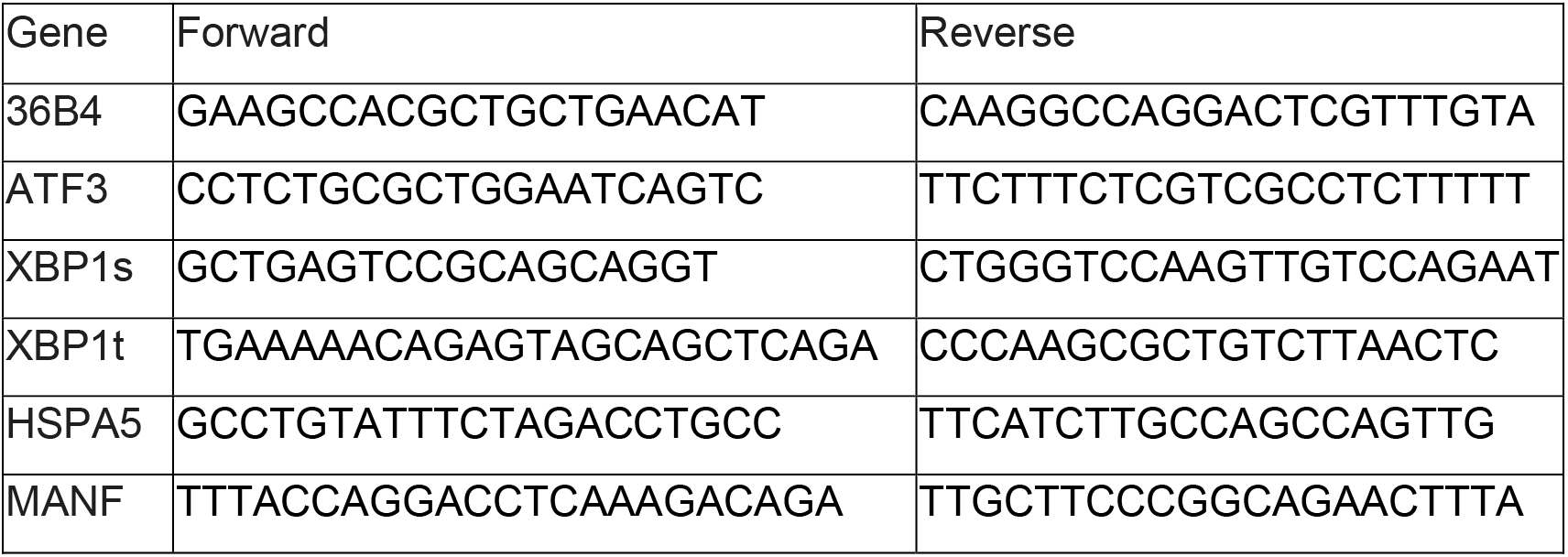

### RNA sequencing analysis

Whole transcriptome RNA sequencing was conducted at the Scripps Research Institute Genomics Core. Libraries were prepared using the NEB Ultra II and then sequenced on an Illumina NextSeq 2000 platform to a depth of 20M 75 bp SE reads per condition. Reads were aligned to the human genome GRCh38 assembly. Differential expression and statistical significance calculations between conditions were assessed with DESeq2 v.1.34.0 in R. Functional gene set enrichment analysis (fGSEA) was performed using the fGSEA package v.s1.20.0 in R. The Hallmark gene set (v7.5.1) was downloaded from MsigDB. UPR gene sets analysis were previously established in (38). Enrichplot package v. 1.14.2 was used to generate the gene set network map.

Single nucleus RNA sequencing was performed at the Scripps Research Institute Genomics Core using 10X Genomics Chromium 3’ v. 3 library preparation protocol and then sequenced on an Illumina NextSeq 2000. Two replicates were performed per condition. FASTQ files were aligned to GRCH38, introns included, using 10X Genomics CellRanger v. 6.0. Quality control metrics were assessed (UMI number, gene number, and mitochondrial percentage). Cells outside a defined range of feature counts and mitochondrial percentages were excluded from downstream analysis (see code for details). Seurat v. 4.0 Canonical Correlation Analysis was used for cluster generation. Harmony v. 0.1 was used for refining sample integration, after which UMAP embedding was performed. Cluster cell identification was performed using markers previously identified in Thomas et al. (21). Progenitor or immature classes were excluded from downstream analyses. Cluster-based fGSEA was performed on pseudobulk data from clusters with DESeq2 calculations for differentially expressed genes. Escape 1.0 was used for single cell fGSEA. The complete RNA-seq data is deposited in gene expression omnibus (GEO) as GSE213948.

## DATA AVAILABILITY STATEMENT

All bulk RNAseq and scRNAseq raw data are deposited in gene expression omnibus (GEO) as GSE213948. Any additional data are available from the authors upon reasonable request.

## ACKNOWLEDGEMENTS

We would like to thank the Lowy Family for their funding support of the MacTel project, the Lowy Medical Research Institute, and this study. Other funding was provided by the National Institutes of Health (AG046495, RFNS125674 to RLW). We would like to thank L. Scheppke for her help editing the manuscript. We would like to thank J. Shimashita and S. Head at the TSRI DNA Array Core for their work on RNA sequencing. We would like to thank J. Orozco for administrative assistance.

---

**Figure S1.**
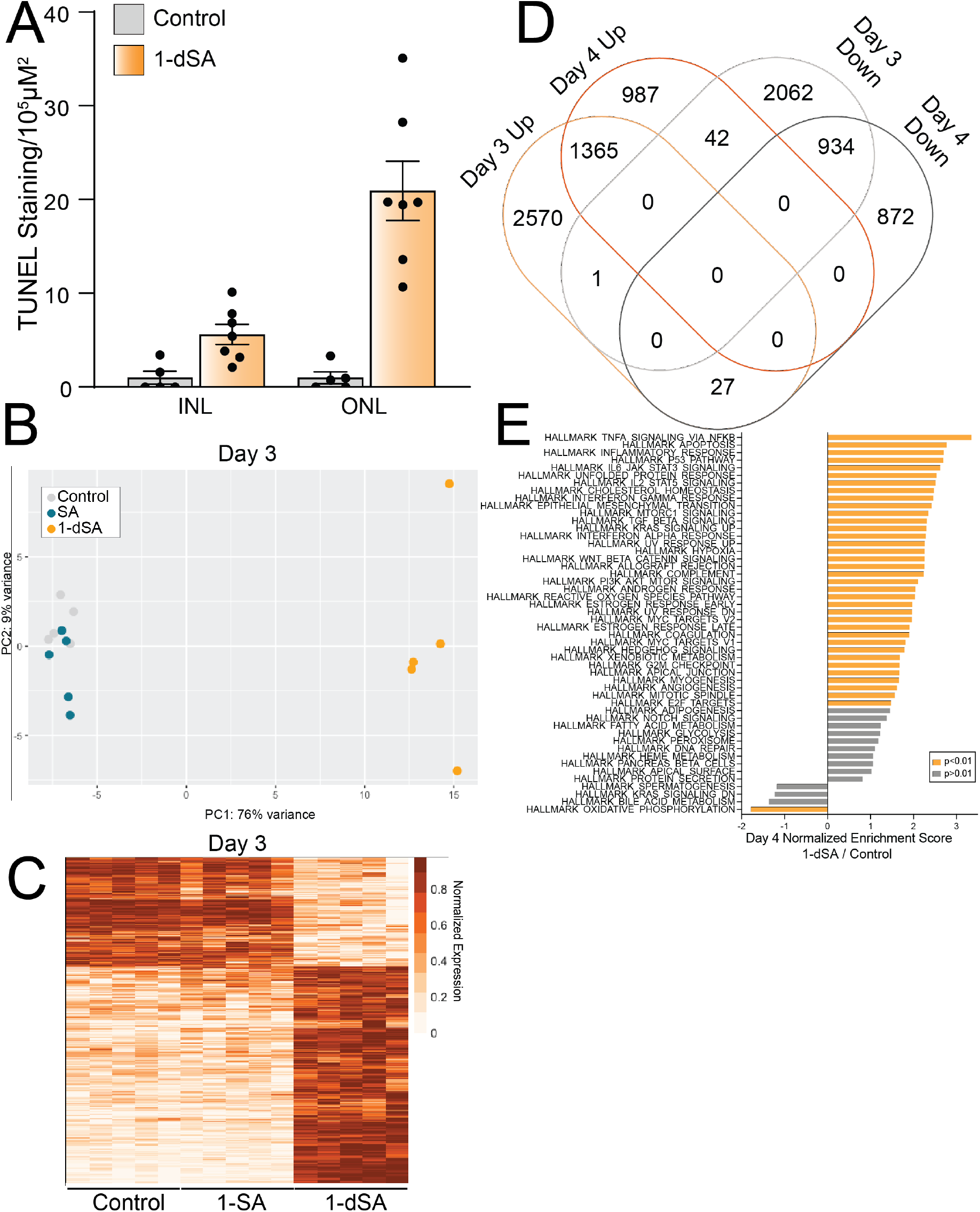
1-dSA-induced transcriptional remodeling precedes cell death. **A**. Quantification of TUNEL staining of ROs treated with 1-dSA (1μM) for 4 days in the inner nuclear layer (INL) or outer nuclear layer (ONL). **B**. Principal component (PC) analysis of all genes from bulk RNAseq data between ROs treated for 3 days with vehicle, sphinganine (SA) (1μM), or 1-deoxysphinganine (1-dSA)(1μM). **C**. Heatmap comparison from bulk RNA-seq between ROs treated for 3 days with vehicle, sphinganine (SA) (1μM), or 1-deoxysphinganine (1-dSA)(1μM) for all genes with p-adj < 0.05 between 1-dSA and vehicle datasets. **D**. Venn diagram showing the overlap of genes increased or decreased in ROs treated for 3 or 4 days with 1-dSA (1 μM), as compared to control. **E**. Enrichment of MSigDB Hallmark pathways in RNAseq data of ROs treated with 1-dSA relative to SA for 4 days. Pathways with enrichment of p-adj < 0.01 are highlighted in orange.

**Figure S2.**
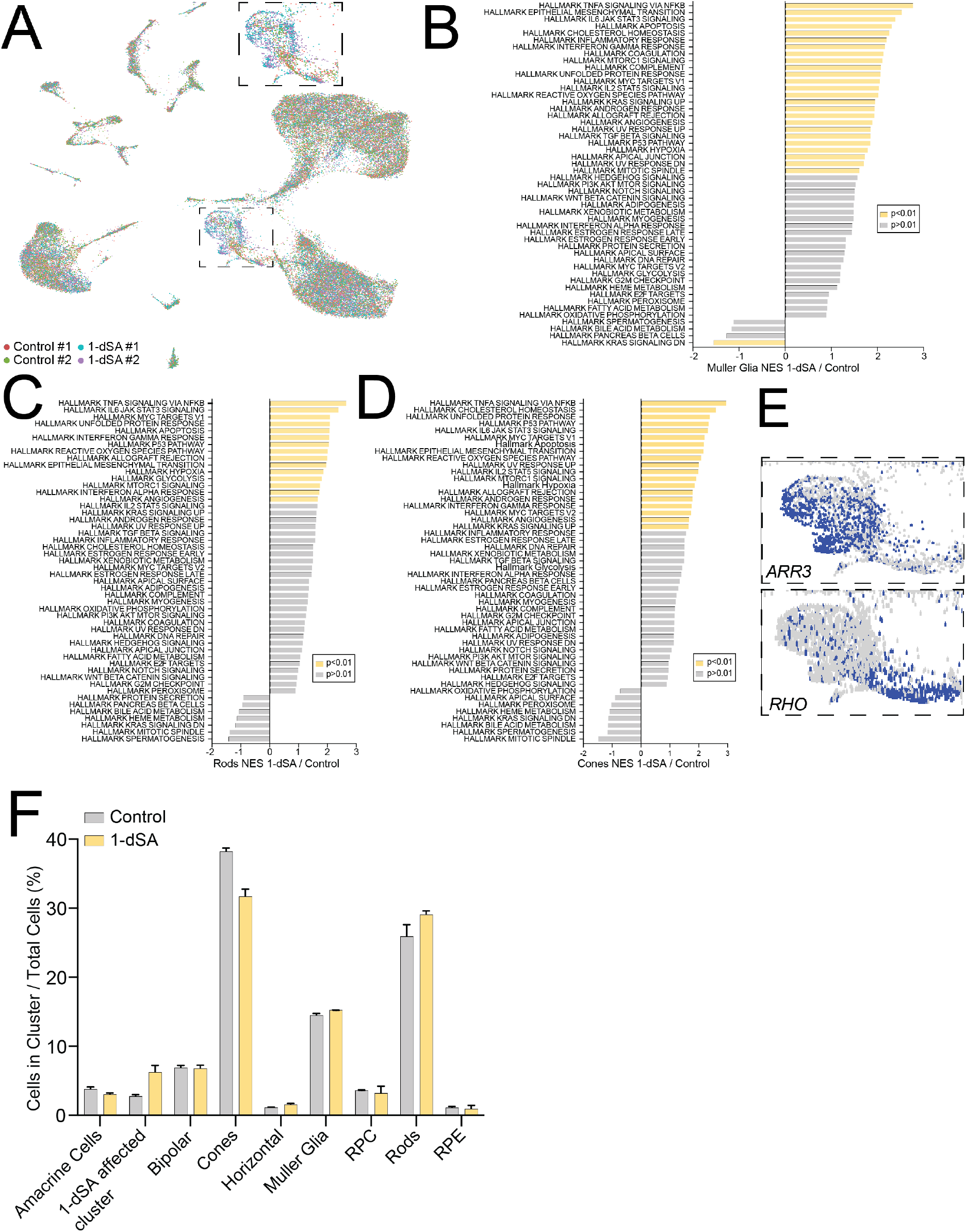
Photoreceptors and Müller glia show a transcriptional response to 1-dSA. **A**. UMAP of unbiased clustering of snRNAseq data from ROs treated with control or 1-dSA (1 μM) for 3 days colored by sample. **B-D**. Enrichment of MSigDB Hallmark pathways in the Müller glia cluster (**B**), rod cell cluster (**C**), and cone cell cluster (**D**) in control organoids relative to 1-dSA treated organoids (1 μM, 3 days). Pathways with enrichment of p-adj < 0.01 are highlighted in yellow. **E**. Expression of the cone marker *ARR3* and the rod marker *RHO* in the 1-dSA affected cluster. **F**. Relative recovery of individual cell types from ROs treated for 3 days with 1-dSA (1 μM, yellow) relative to control (grey).

**Figure S3.**
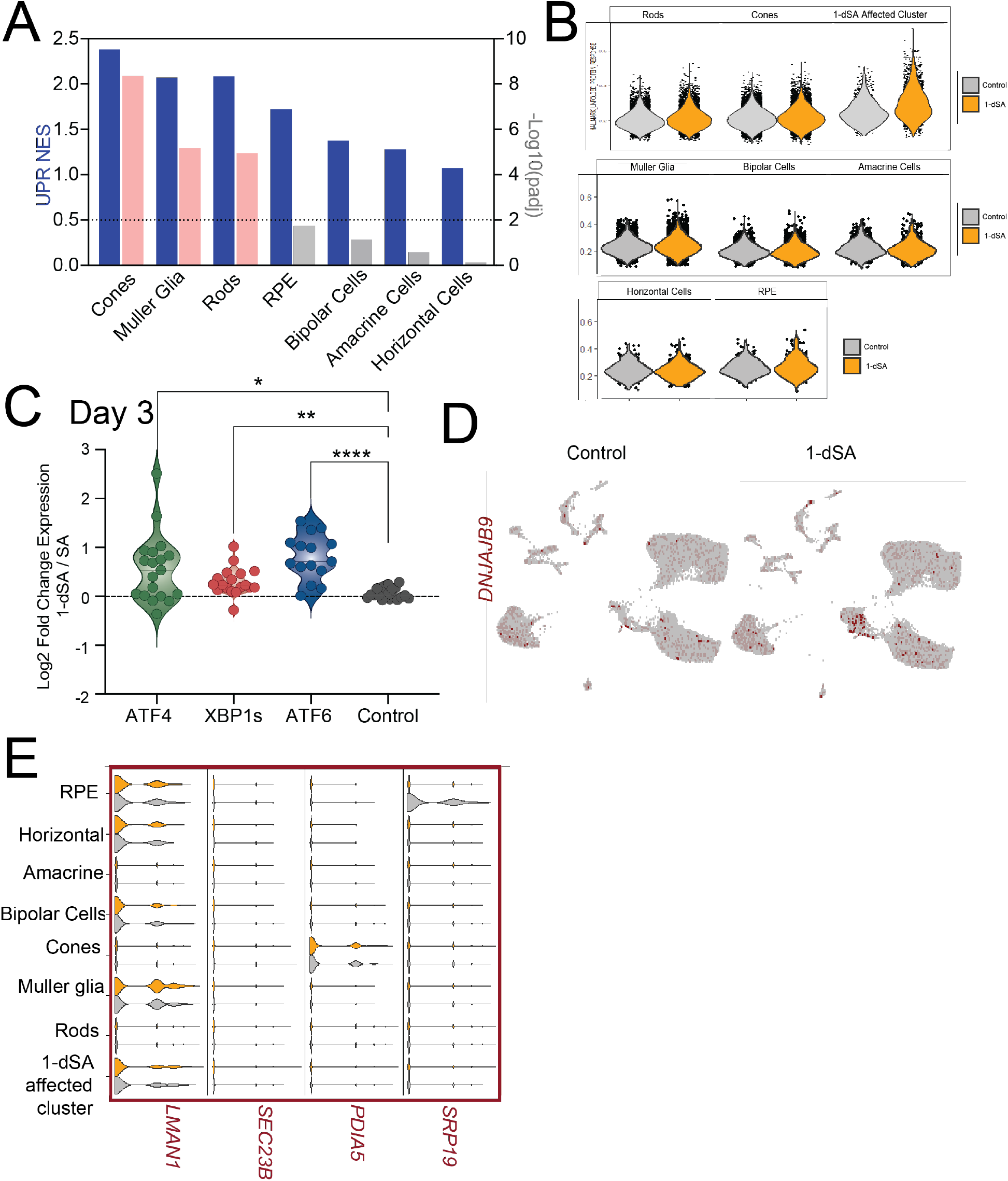
The UPR is differentially activated in photoreceptors and Muller glia from 1-dSA treated organoids. **A**. MSigDB Hallmark fGSEA analysis for Unfolded Protein Response (UPR). Normalized enrichment scores (NES, blue) and −log10(pink: p-adj <0.01; grey: p-adj > 0.01) for 1-dSA treated cells relative to control cells from snRNAseq dataset are shown. **B**. Escape single-cell analysis of MSigDB Unfolded Protein Response (UPR) enrichment per cell in snRNAseq dataset from control (grey) or 1-dSA (1 μM, 3 days) treated organoids. **C**. Quantification of RNAseq fold changes of gene targets of the UPR-induced transcription factors ATF4, XBP1s, and ATF6 in ROs treated for 3 days with 1-dSA (1 μM) relative to those treated with sphinganine (SA, 1 μM). *p<0.05, **p<0.01, ***p<0.005 for Brown-Forsythe and Welch ANOVA tests compared to control gene set. **D**. Expression of the XBP1s target gene *DNAJB9* across cell types in control or 1-dSA (1 μM, 3 days) treated ROs. **E**. Violin plot of gene targets of XBP1s target genes from our snRNAseq dataset separated by cluster and treatment. 1-dSA-treated organoids (1 μM, 3 days) are in orange; control-treated organoids are in grey.

**Figure S4.**
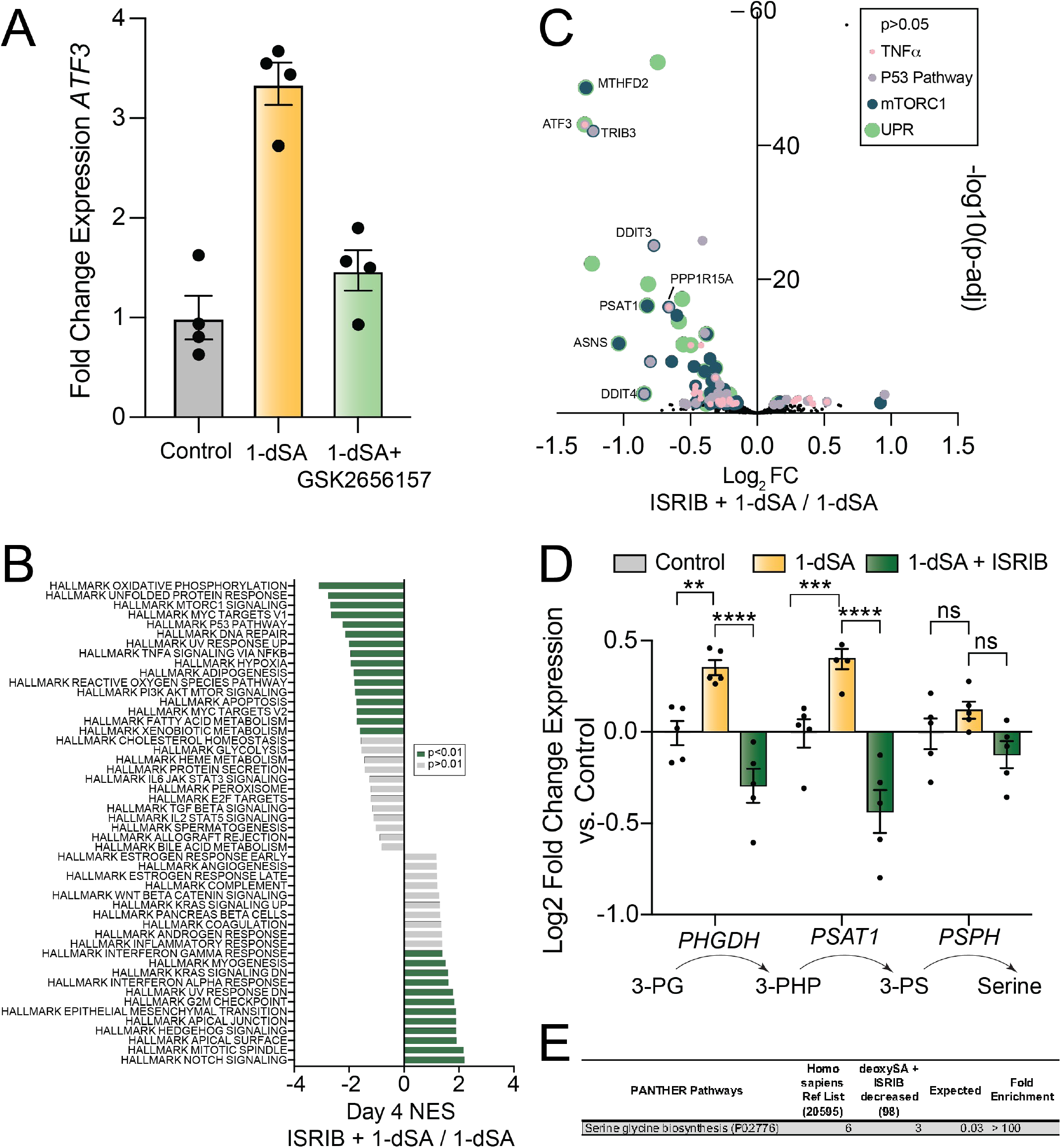
PERK/ISR signaling contributes to 1-dSA-induced retinal toxicity. **A**. qPCR of the ATF4 gene target *ATF3* in ROs treated with vehicle, 1-dSA, or 1-dSA and GSK2656157. **B**. Enrichment of MSigDB Hallmark pathways in the Müller glia cluster in 1-dSA- and ISRIB-treated organoids relative to 1-dSA-treated organoids. Pathways with enrichment of p-adj < 0.01 are highlighted in green. **C**. Scatterplot of differentially expressed MSigDB pathway genes, as labeled, in ROs treated with ISRIB and 1-dSA relative to 1-dSA. Pathways that contain the same genes show overlapping markers. **D**. Expression fold changes of the serine synthesis genes *PHGDH, PSAT1*, and *PSPH* from bulk RNA-seq data of ROs treated with 1-dSA (1μM) in the presence or absence of ISRIB for 4 days compared to control-treated ROs. **p<0.01, *** p<0.001, ****p<0.0001 for two-way ANOVA. **E**. Gene ontology analysis of panther pathways for differentially expressed genes with p-adj < 0.05 and Log2FC < −0.5 between ROs treated with ISRIB and 1-dSA relative to 1-dSA-treated ROs.

**Figure S5.**
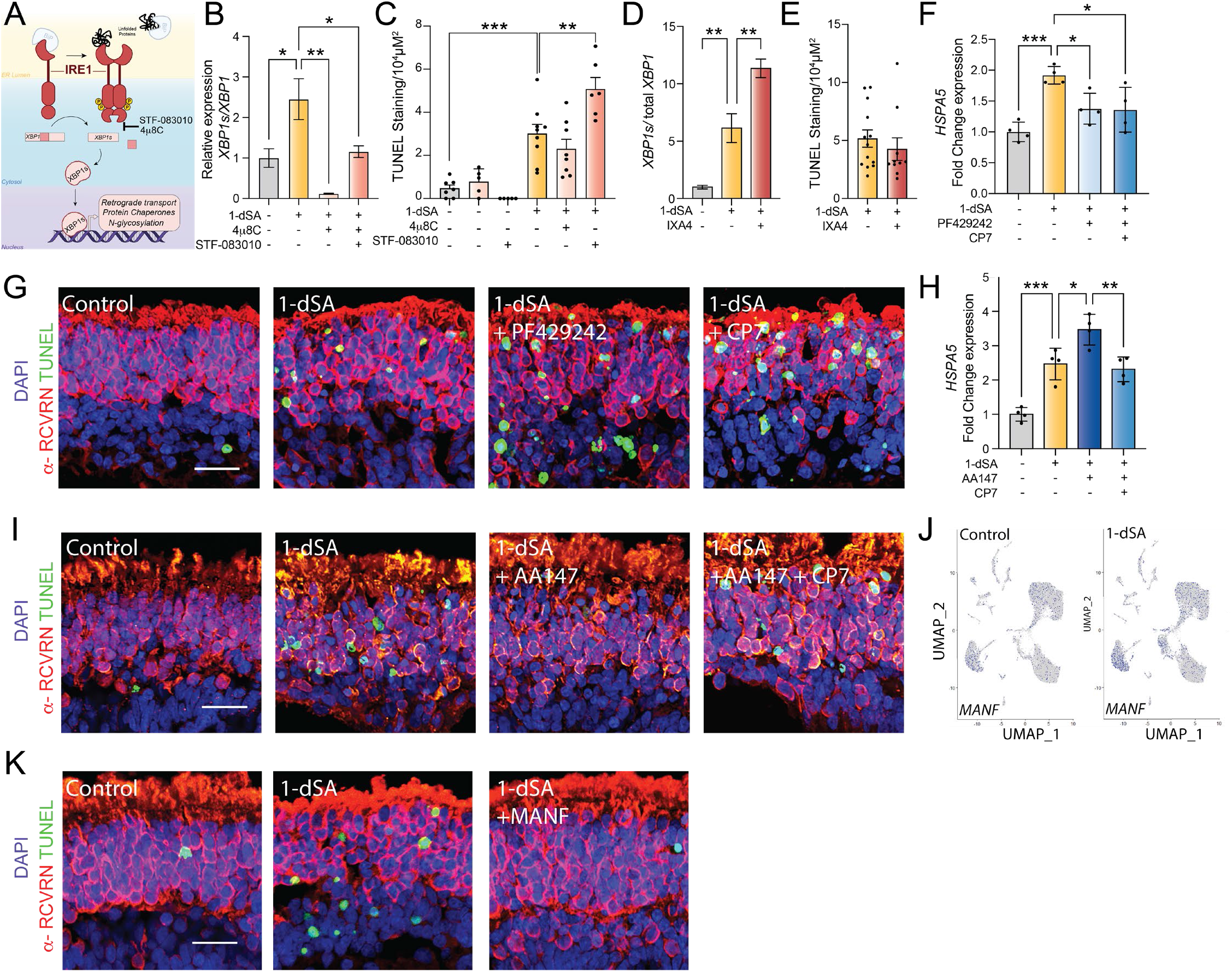
ATF6 activity protects the retinal from 1-dSA toxicity. **A**. Illustration of the IRE1/XBP1s UPR signaling pathway and pharmacologic inhibitors of this pathway. **B**. qPCR measuring expression of spliced *XBP1 (XBP1s*) relative to total *XBP1* in ROs treated for 4 days with 1-dSA in the presence or absence of the IRE1 inhibitors 4μ8C (32 μM) or STF-083010 (32 μM). *p<0.05, **p<0.01 for ordinary one-way ANOVA. **C**. Quantification of TUNEL staining of ROs treated with 1-dSA (1μM) for four days in the presence or absence of the IRE1 inhibitors 4μ8C (32 μM) or STF-083010 (32 μM). **p<0.01, ***p<0.001 for ordinary one-way ANOVA. **D.** qPCR of the expression of spliced *XBP1* relative to total *XBP1* in ROs treated for 4 days with 1-dSA (1 μM) with or without the IRE1 activator IXA4 (10 μM). **p<0.01 for ordinary one-way ANOVA. **E**. Quantification of TUNEL staining of ROs treated with 1-dSA (1μM) for four days in the presence or absence of the IRE1 activating compound IXA4 (10 μM). **F**. qPCR of the ATF6 gene target *HSPA5* in ROs treated with 1-dSA (1 μM) and Ceapin-A7 (CP7; 7 μM) or PF429242 (10 μM). *p<0.05, **p<0.01, ***p<0.001 for ordinary one-way ANOVA. **G**. Representative images of retinal organoids treated with 1-dSA (1 μM), Ceapin-A7 (CP7; 7 μM), or PF-429242 (10 μM) for 4 days. DAPI staining, TUNEL staining, and α-recoverin (α-RCVRN) staining are shown. **H**. qPCR of the ATF6 gene target *HSPA5* in ROs treated with 1-dSA (1 μM) in the presence or absence of AA147 (10 μM) and Ceapin-A7 (CP7; 7 μM). *p<0.05, **p<0.01, ***p<0.001 for ordinary one-way ANOVA. **I**. Representative images of retinal organoids treated with 1-dSA (1 μM), AA147 (10 μM), or Ceapin-A7 (CP7; 7 μM) for 4 days. DAPI staining, TUNEL staining, and α-recoverin (α-RCVRN) staining are shown. **H**. Expression of *MANF* across cell types in ROs treated with 1-dSA (1 μM; 3 days). **K**. Representative images of retinal organoids treated with 1-dSA (1 μM), and MANF (100 ng/μL) for 4 days. DAPI staining, TUNEL staining, and α-recoverin (α-RCVRN) staining are shown. Scale bar is 25μm.

## SUPPLEMENTAL TABLE LEGENDS (see included Excel sheets for the Tables)

**Supplementary Table S1. RNAseq of human iPSC-derived retinal organoids treated with 1-dSA for 2 days**. The included excel spread shows the DESEQ2 analysis of RNAseq of retinal organoids treated with vehicle or 1-dSA (1 μM) for 2 days. The complete RNAseq data is deposited in gene expression omnibus (GEO) as GSE213948.

**Supplementary Table S2. RNAseq of human iPSC-derived retinal organoids treated with 1-SA or 1-dSA for 3 days**. The included excel spread shows the DESEQ2 comparisons of RNAseq data from human iPSC-derived retinal organoids treated with 1-SA (1 μM), 1-dSA (1 μM), or vehicle equivalent for 3 days. The complete RNAseq data is deposited in gene expression omnibus (GEO) as GSE213948.

**Supplementary Table S3. RNAseq of human iPSC-derived retinal organoids treated with 1-dSA for 4 days**. The included excel spread shows the DESEQ2 comparisons of RNAseq data from human iPSC-derived retinal organoids treated with 1-dSA (1 μM) or vehicle equivalent. The complete RNAseq data is deposited in gene expression omnibus (GEO) as GSE213948.

**Supplementary Table S4. snRNAseq of human iPSC-derived retinal organoids treated with 1-dSA for 3 days**. The included excel spread shows the DESEQ2 comparisons of snRNAseq data from human iPSC-derived retinal organoids treated with 1-dSA (1 μM) relative to vehicle equivalent. Clusters are labeled as noted in **Fig. 2A**. The complete RNAseq data is deposited in gene expression omnibus (GEO) as GSE213948.

**Supplementary Table S5. RNAseq of human iPSC-derived retinal organoids treated with 1-dSA for 4 days in the presence or absence of ISRIB**. The included excel spread shows the DESEQ2 comparisons of RNAseq data from human iPSC-derived retinal organoids treated with 1-dSA (1 μM) in the presence or absence of ISRIB (200 nM). Comparisons are as noted in the excel file. The complete RNAseq data is deposited in gene expression omnibus (GEO) as GSE213948.

**Supplementary Table S6. RNAseq of human iPSC-derived retinal organoids treated with 1-dSA for 4 days in the presence or absence of AA147**. The included excel spread shows the DESEQ2 comparisons of RNAseq data from human iPSC-derived retinal organoids treated with 1-dSA (1 μM) in the presence or absence of AA147 (10 μM). Comparisons are as noted in the excel file. The complete RNAseq data is deposited in gene expression omnibus (GEO) as GSE213948.

